# Acute Exposure to Perfluorooctanoic Acid (PFOA) During Cardiomyogenesis disrupts Transcriptional and Electrophysiological Profiles in Differentiated Myocytes

**DOI:** 10.64898/2026.05.05.723050

**Authors:** Tomoko Ishikawa, Christopher W. Clark, Anagha Tapaswi, Kimberley E. Sala-Hamrick, Todd J. Herron, Eric N. Jimenez Vazquez, Abhilasha Jain, David K. Jones, Justin A. Colacino, Andre Monteiro Da Rocha, Laurie K. Svoboda

**Affiliations:** Department of Environmental Health Sciences, School of Public Health, University of Michigan, Ann Arbor, MI 48109, USA; Department of Internal Medicine, Medical School, University of Michigan, Ann Arbor, MI 48109, USA; Greenstone Biosciences, Palo Alto, CA, 94304, USA; Department of Pharmacology, Medical School, University of Michigan, Ann Arbor, MI 48109, USA; Department of Nutritional Sciences, School of Public Health, University of Michigan, Ann Arbor, MI 48109, USA

**Keywords:** PFAS, DOHaD, induced pluripotent stem cell, optical mapping, cardiomyocyte

## Abstract

The early developmental environment plays a critical role in the etiology of cardiovascular diseases (CVDs), but underlying molecular mechanisms are poorly understood. Exposure to per and polyfluoroalkyl substances (PFAS) are linked to various CVDs, but effects of developmental PFAS exposures on the human heart remain unclear. Using human-induced pluripotent stem cell-derived cardiomyocytes (hiPSC-CM), the objective of this study was to investigate the effects of PFAS exposure during cardiac differentiation on gene expression and function of cardiomyocytes. We exposed two hiPSC lines (one male and one female donor) to perfluorooctanoic acid (PFOA), a common and ubiquitous PFAS (0.05, 0.5, 5, 50, 100, 150, 200 μM), followed by assessment of cellular number and pluripotency marker expression. PFOA exposure for 72 hours had no significant effects on hiPSC pluripotency, and modest inhibition of proliferation was observed only at the highest concentration. hiPSCs were then differentiated into ventricular cardiomyocytes in the continued presence or absence of PFOA (0, 0.5, 5, 50 μM) using an established small molecules protocol. Optical mapping studies using voltage and calcium-sensitive dyes revealed dose and cell line-specific effects of PFOA on cardiomyocyte voltage and calcium dynamics that were still present 10 days after cessation of exposure. Patch clamping studies demonstrated small but significant reductions in repolarizing *I_Kr_* currents with 5µM PFOA exposure in cardiomyocytes from both donors. Using RNA-seq, we found that exposure to PFOA led to significant changes in transcriptional pathways related to lipids and lipoproteins in the female hiPSC-CM. In the male hiPSC-CM, we observed significant effects on developmental pathways and calcium homeostasis. Thus, we found that environmentally relevant PFOA exposure during cardiomyocyte differentiation affects the electrophysiological properties and transcriptome of hiPSC-CM even after cessation of exposure, with effects that differ by donor cell line. These findings provide direct experimental evidence that transient developmental exposure to PFOA can durably reprogram human cardiomyocyte function, supporting a developmental origin of PFAS-associated cardiovascular risk.

**Impact Statement:** These studies demonstrate that exposure to environmentally relevant levels of PFOA during the differentiation of hiPSCs into cardiomyocytes alters cardiac gene expression and function, with effects that persist beyond cessation of exposure.

## Introduction

Per- and poly-fluoroalkyl substances (PFAS) are a group of manmade chemicals used in a wide array of consumer products^1^. Structurally, PFAS consist of a chain of carbon atoms bound to fluorine atoms, with a functional group at the end of the molecule. PFAS can vary in chain length, as well as in functional groups and degree of branching. The carbon-fluorine bonds in PFAS make them highly chemically stable, bioaccumulative, and resistant to environmental degradation. Widespread global use of PFAS, combined with their environmental persistence, has resulted in contamination of indoor and outdoor spaces^2,3^, drinking water^4,5^, and food^6–8^ worldwide. Human exposure is ubiquitous, and PFAS have been detected in the blood of almost every human being on the planet^9,10^. The Developmental Origins of Health and Disease (DOHaD) hypothesis states that exposure to environmental factors such as PFAS during early development may have long-term, adverse health consequences. PFAS exposures clearly occur during development, as PFAS have been detected in the placenta and fetus, as well as in breast milk^5,11–13^. Exposure to PFAS in adults is associated with several cardiovascular diseases, including atherosclerosis, ischemic heart disease, and myocardial infarction^14–17^. Less is known, however, about the cardiovascular effects of PFAS exposures that occur early in life. Human ^18,19^ and animal ^20,21^ studies show that PFAS exposures during development can have adverse cardiovascular effects, including congenital defects in cardiac function and morphology; however, it remains unknown whether transient developmental PFAS exposure induces persistent functional reprogramming in human cardiomyocytes.

In this study, we utilized human induced pluripotent stem cells (hiPSCs), a new approach methodology, to investigate the effects of PFAS exposure during differentiation on cardiomyocyte function. The hiPSC-cardiomyocyte model recapitulates many aspects of human embryonic heart development^22^, providing genetic tractability and temporal resolution that are not available in human population or animal studies. We directed cells to undergo differentiation to ventricular cardiomyocytes using a published GiWi protocol^23^ with minor modifications^24^ and exposed the cells to perfluorooctanoic acid (PFOA), a common and ubiquitous legacy PFAS that is still widespread in the environment and in human blood. We then examined the effects of exposure to PFOA on cardiomyocyte electrophysiology and gene expression. Our findings demonstrate that exposure to this persistent organic pollutant during human cardiomyocyte differentiation has effects on cardiomyocyte gene expression and electrophysiology that persist after cessation of exposure.

## MATERIALS AND METHODS

### hiPSC culture and cardiomyocyte differentiation

hiPSC lines derived from one male (PENN042i-258-12) and one female (PENN002i-442-1) donor were purchased from WiCell Research Institute (Madison, Wisconsin). Cells were seeded on Matrigel-coated plates (Corning), maintained with iPS-Brew XF (StemMACS), and split once a week using Versene (Gibco). The hiPSCs were replated as a monolayer for directed differentiation into ventricular cardiomyocytes using an established GiWi protocol with minor modifications^23,24^. When the monolayer reached 80-90% confluency, differentiation was initiated (day 0) by applying 4 μM CHIR 99021 (Tocris) in RPMI-1640 media supplemented with 0.5% bovine serum albumin (Sigma) and 0.2% ascorbic acid (Sigma). After 48 hours (day 2), 5 μM IWP4 (Tocris) was applied to promote the induction of cardiac mesoderm. After D8, the cells were kept in maintenance medium (RPMI-1640 supplemented with B27). Twenty days after the initiation of the differentiation, the cardiomyocytes were purified using a magnetic separation system with a PSC-Derived Cardiomyocyte Isolation Kit (Miltenyi Biotec, Gaithersburg, MD) and replated with EB20 (DMEM:F12 supplemented with 10% FBS, 1 mmol/L L-glutamine, 0.1 mmol/L nonessential amino acids, 0.1 mmol/L β-mercaptoethanol and 25µM of blebbistatin) on CELLvo^TM^ Matrix Plus plates (StemBioSys) without any PFOA. Matrix Plus plates promote maturation of hiPSC-CMs to a more adult-like morphology and function^25^. Two days after purification, the media was changed to maintenance medium without any PFOA for 8 additional days. All cells were maintained in an incubator at 37°C with 5% CO_2_ in atmospheric air.

### Cell line genotyping for sex

To verify the sex of the patient donors, we conducted PCR for expression of *SRY* (expressed on the Y-chromosome) and *ATL1* (expressed on the X-chromosome). After several passages, hiPSCs from each patient donor were collected, and DNA was extracted using the AllPrep DNA/RNA/Protein kit (Qiagen). DNA was amplified with HotStarTaq Master Mix kit (Qiagen) and published^26^ primer set *SRY* (F: 5’-CATGAACGCATTCATCGTGTGGTC-3’ and R: 5’-CTGCGG GAAGCAAACTGCAATTCTT-3’ ) and *ATL1* (F: 5’-CCCTGATGAAGAACTTGTATCTC-3’ and R: 5’-GAAATTACACACATAGGTGGCACT-3’). The PCR reaction was performed using the C1000 Touch Thermal Cycler (Bio-Rad) at 95°C (15 min) for initial DNA denaturation, followed by 35 cycles of 94°C (30 sec), 62°C (30 sec), 72°C (1 min) with a final extension at 72°C (10 min). PCR products and size markers were run on the QIAxcel (Qiagen). We confirmed expression of *SRY* only in the male hiPSC line, and expression of *ATL1* at a level approximately twice as high in the female line compared to the male line, as expected (Supplemental Figure 1).

### Time-lapse imaging of cellular confluence

hiPSCs were seeded into 24-well plates and cultured for 48 hours before being exposed to PFOA for 72 hours. Cellular confluence, i.e. the total area of the plate covered by hiPSC colonies, was monitored using the IncuCyte ZOOM live cell imaging system (Essen BioScience) with bright field image acquisition every 6 hours and quantified using IncuCyte Zoom 2018A software.

### Assessment of hiPSC pluripotency

hiPSCs were collected for RNA extraction at 55-70% confluence. RNA was extracted using the AllPrep DNA/RNA/Protein kit (Qiagen) following the manufacturer protocol. RNA was reverse transcribed into cDNA using the iScript cDNA synthesis kit (Bio-Rad). Gene expression of the pluripotency markers, *POU5F1 (OCT4)*, *SOX2*, and *NANOG* was assessed using RT-qPCR (CFX384 Touch Real-time PCR Detection System, Bio-Rad) with SeoAdvanced Universal SYBR Green Supermix (Bio-Rad). Relative quantification was performed using the geometric mean of three housekeeping genes (*GAPDH*, *ATCB*, and *RPS18*) in each experiment. All primers (PrimePCR^TM^ SYBR® Green Assay) were purchased from Bio-Rad. Catalog numbers were as follows: *POU5F1* (qHsaCED0038334), *SOX2* (qHsaCED0036871), *NANOG* (qHsaCED0043394), *GAPDH* (qHsaCED0038674), *ATCB* (qHsaCED0036269), and *RPS18* (qHsaCED0037454).

### PFOA Exposure

PFOA (Cat# 171468) was purchased from Sigma (St. Louis, MO). Working solutions of PFOA (0.5-50 mM) were prepared in molecular-grade dimethyl sulfoxide (DMSO) (Sigma) and added to the media at a 1:1000 ratio before every use. On the first day of exposure, hiPSCs were exposed to the DMSO vehicle or each concentration of PFOA (0, 0.5, 5, and 50µM) in Hanks’ Balanced Salt Solution without phenol red (HBSS, Gibco) for 1 hour. HBSS was then removed, and media with each concentration of PFOA was added. Medium containing PFOA or vehicle was replaced daily. After 72 hours of exposure, hiPSCs were dissociated using Versene (Gibco) and re-plated for differentiation as described above. During differentiation, PFOA exposure was refreshed with each media change and continued until day 20 after initiation of differentiations. The paradigm for differentiation and PFOA exposures is shown in **Figure 1**. Biological replicates were defined as experiments on separate days using different cell passages. We performed 3 biological replicates, each with 4-6 technical replicates, per condition.

**Figure 1:**
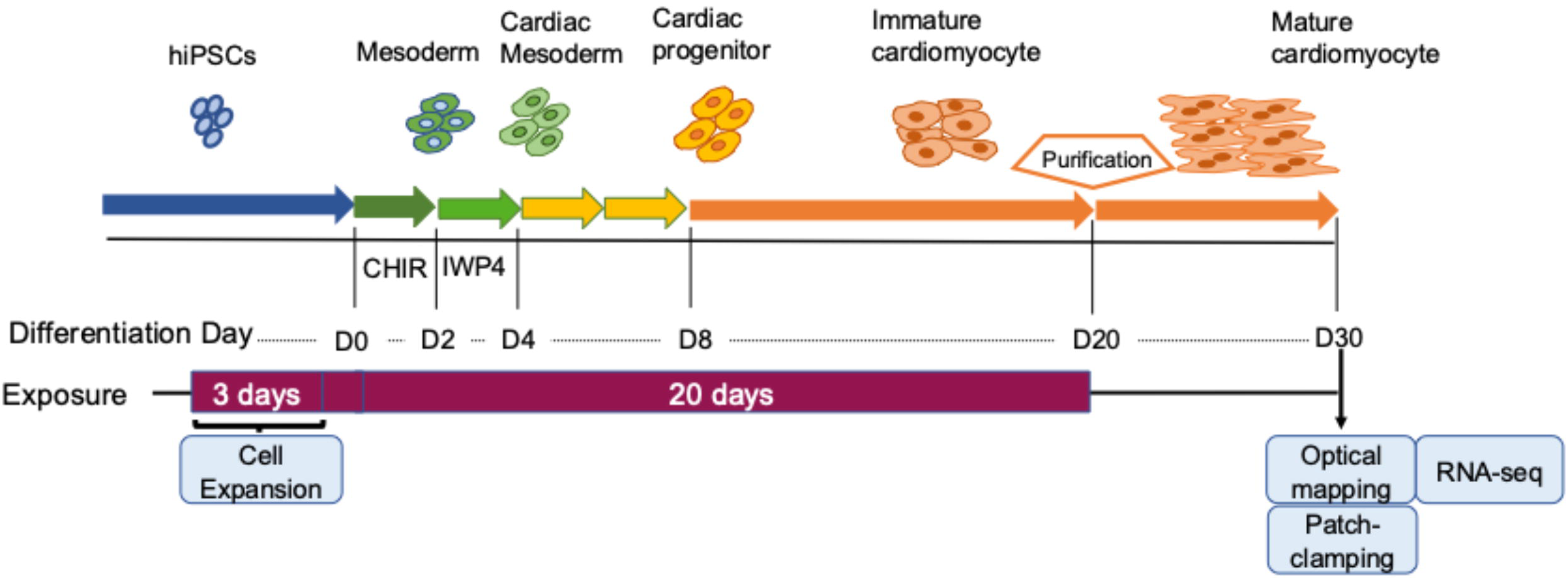
Paradigm for hiPSC differentiation, PFOA exposure, and experimental endpoints.

### Cardiomyocyte Purification and Optical Mapping

Cardiomyocyte purification was performed using a magnetic-activated cell separation system with the PSC-Derived Cardiomyocyte Isolation Kit (Miltenyi Biotec) following the manufacturer protocol, with minor modifications^24^. Purified cardiomyocytes were seeded in 96-well CELLvo Matrix Plus plates (StemBiosys) at a density of 7.5×10^4^ cells/well with EB20 media and maintained with B27(-) media for ten days to allow for cardiomyocyte maturation^25^. Action potentials and calcium transients were recorded using the FluoVolt membrane potential probe (Life Technologies), and Calbryte 520 AM, intracellular free-Ca probe (AAT Bioquest), respectively. After the cells were rinsed with HBSS, they were incubated with either probe for 30 minutes at 37°C, rinsed again, then placed back in HBSS. When the HBSS reached 37°C, fluorescence signal images were recorded for 30 seconds at a frame rate of 250 frames per second and LED intensity of 10.0% for action potentials and 5.0% for calcium transients using the Nautilus high-throughput optical mapping system (Curi Bio). The fluorescence signal images were converted to electrophysiological parameters using the StemBioSys Optical Electrophysiology Analysis Tool software (StemBiosys). Action potential and calcium transient duration data were corrected for beat frequency using Fridericia’s equation: 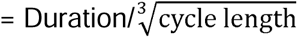. Cycle length = 1/beat frequency in Hz. After the recordings were completed, the plates were then rinsed with HBSS and stored at -80°C for molecular analyses.

### Patch Clamp

After 10 days of maturation on the Matrix Plus plates, the cardiomyocytes were re-plated onto Matrigel-coated cover glasses for patch clamp. Patch clamping was conducted as previously described^27^. Standard patch clamp techniques were used to measure *I_Kr_*, *I_K1_*, and *I_Ca_*. All measurements were conducted at 37°C using a whole-cell patch clamp equipped with an IPA Integrated Patch Amplifier acquired with SutterPatch (Sutter Instrument) and analyzed with Igor Pro 8 software (Wavemetrics). Data were sampled at 5 kHz and processed with a low-pass filter at 1 kHz. The cells were perfused in extracellular solution containing (in mM): 150 NaCl, 5.4 KCl, 1.8 CaCl_2_, 1 MgCl_2_, 15 Glucose, 10 HEPES, 1 Na-Pyruvate and titrated to pH 7.4 using NaOH. Recording pipettes had resistances between 2 to 5 MΩ when backfilled with intracellular solution containing (in mM): 5 NaCl, 150 KCl, 2 CaCl2, 5 EGTA, 10 HEPES, 5 MgATP and titrated to pH 7.2 using KOH. We stored single-use 1 mL aliquots of intracellular solution at - 20°C until the day of recording and kept thawed aliquots on ice throughout all recordings. Sodium currents were inactivated by applying a 100 ms step to -40 mV before recording *I_K1_*and *I_Kr_*. To activate *I_hERG_*, we stepped cells from a holding potential of −50 mV for 1 s to a 3 s pre-pulse between −50 mV and +50 mV in 10 mV increments. Tail currents were then measured during a −40 mV, 6 s test pulse. We normalized peak tail current to cellular capacitance, plotted current density as a function of pre-pulse potential in mV, and fitted the data with the following Boltzmann equation:

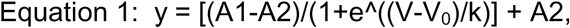

where A1 and A2 represent the maximum and minimums of the fit, respectively, V is the membrane potential, V_0_ is the midpoint, and *k* is the slope factor. The steady-state current was measured as the average current during the last 10 ms of each step pulse.

### RNA Sequencing

RNA was extracted from CMs using the AllPrep DNA/RNA/miRNA Universal kit (Qiagen) following the company’s protocols. Quality control of the cDNA was performed using D5000 ScreenTape at the UM Advanced Genomics Core. Libraries for RNA-seq were prepared using a plexWell kit (seqWell) following a modified version of the manufacturer protocol as previously described^28^. Libraries were sequenced on the NovaSeq X (Illumina) at the UM Advanced Genomics Core. The quality of sequencing reads was assessed using FastQC^29^(v0.12.1), and MultiQC^30^(v1.8). The reads were aligned to the reference human genome (GRCh38) using STAR^31^ with default parameters. Principal components analysis (PCA) was performed with a subset of the top 1,000 variable genes after genes on Chromosome X, Y, and mitochondria were eliminated. Batch effects were adjusted using ComBat-seq^32^. Differentially expressed genes (DEGs) were identified using edgeR^33^. Gene set enrichment analyses for gene ontology were performed using clusterProfiler^34^ gseGO with genome-wide annotation for Human, org.Hs.eg.db. P-values were adjusted using Bonferroni correction, and an FDR <0.05 was considered significant. We also examined selected calcium-handling genes individually. The counts of those genes were extracted from RNA sequence data, and RPKM (Reads Per Kilobase per Million reads) values were calculated by normalizing for sequencing depth and gene length.

### Statistical Analysis

All statistical analysis was conducted in R version 4.4.0. Cell confluence data were analyzed using linear regression within each cell line comparing the growth rate for each PFOA concentration to control. Statistical analysis of optical mapping was performed using one-way ANOVA and followed by post hoc tests using the Bonferroni correction within each cell line. Statistical analysis of the patch clamp data was performed using a non-parametric Mann-Whitney test or two-way ANOVA with the Mann-Whitney test after being tested for normality (Shapiro-Wilk test) and for outlier identification (ROUT and Grubbs’ tests).

## RESULTS

### Effects of PFOA on hiPSC proliferation

Before assessing effects of PFOA on cardiac differentiation, we first examined the potential cytotoxic effects of PFOA exposure in the hiPSCs in response to a range of PFOA concentrations. The concentrations encompass those historically found in human serum in occupational settings (50 μM = 20.7μg/mL, 5 μM = 2.07μg/mL)^35^, those found in highly exposed communities (0.5 μM = 207ng/mL)^4,36^ and supra-physiological concentrations (200 μM). We utilized the IncuCyte system for time-lapse image monitoring of the effects of PFOA exposure on cellular confluence, a read-out of cellular proliferation and viability. hiPSCs were exposed to 0-200 μM PFOA, and the area of the plate covered by cells was monitored between 0 and 72 hours. Growth curves for each PFOA concentration were compared to control using linear regression (Figure 2). Cell proliferation continued across all conditions, with 200 μM PFOA significantly slowing the rate of proliferation in both cell lines. Interestingly, we observed slightly higher proliferation rates at some of the lower PFOA concentrations compared to control. In the female hiPSC line, we found a statistically significant increase in the growth rate at 0.5 μM PFOA (p=0.004). In the male hiPSC line, the increase was significant at the 0.5, 5, and 50 μM concentrations, with 5 μM PFOA exhibiting the highest growth rate (p=0.016, p<0.0001, p=0.002). These findings suggest a potential non-monotonic response, consistent with emerging PFAS toxicity paradigms.

**Figure 2:**
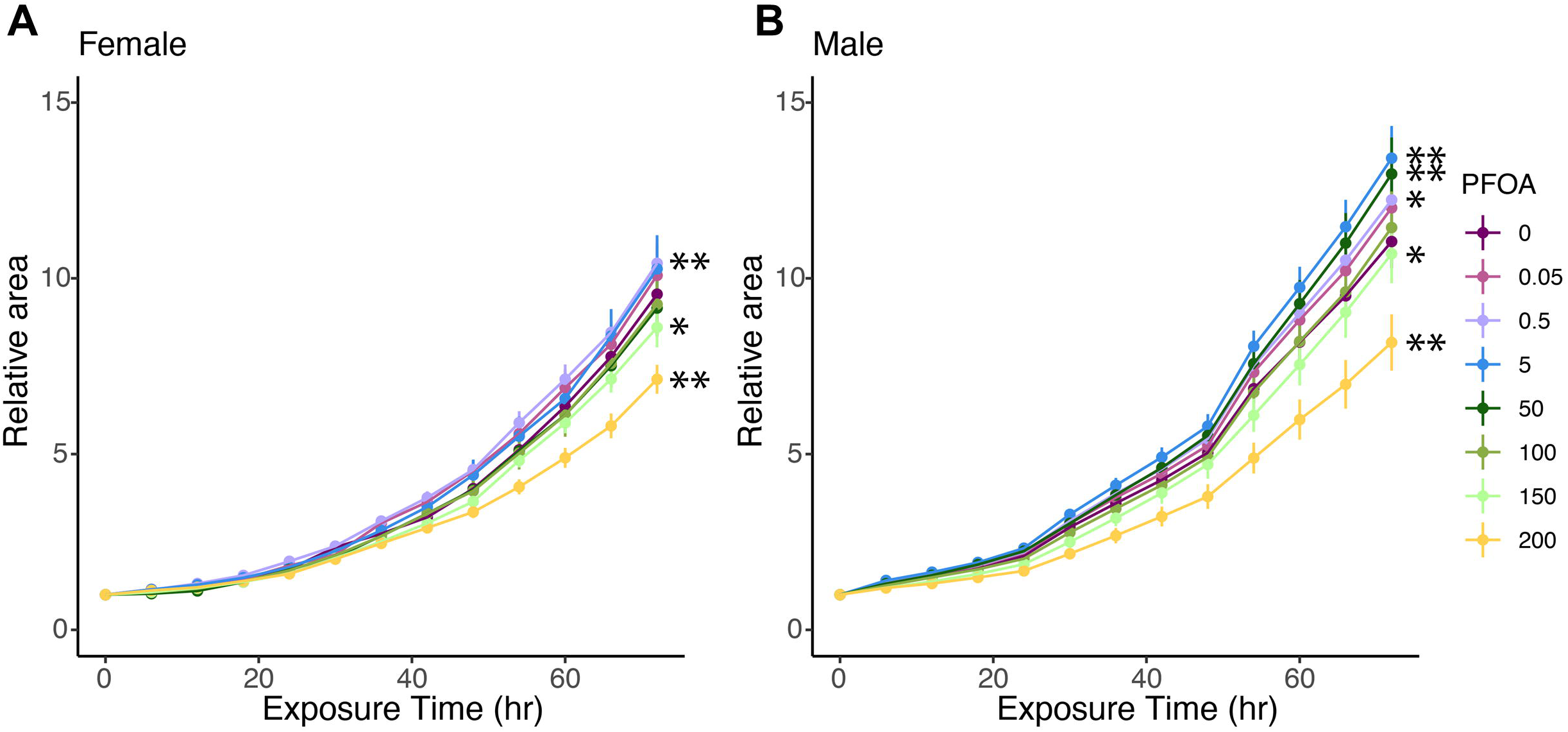
Dose-dependent effects of PFOA exposure on expansion of hiPSCs. Cellular confluence, a readout of proliferation, was monitored in real-time and quantified. Data for each time point are expressed as the fold increase in area covered by cells, relative to time 0. (A) Growth trajectory of the female hiPSC line. (B) Growth trajectory of the male hiPSC line. Data are shown as the mean ± SEM. *p<0.05, **p<0.01.

### PFOA exposure and hiPSC pluripotency

We next examined whether PFOA exposure would impact pluripotency of the hiPSC. Using qRT-PCR, we assessed the effects of 72-hour PFOA exposure on gene expression of the pluripotency markers, *POU5F1 (OCT4)*, *SOX2*, and *NANOG*. PFOA had no significant effects at any concentration on the expression of these genes in either cell line. Notably, baseline (control) expression of *NANOG* was consistently lower in the male hiPSC line compared to the female line, and this difference was statistically significant in the control group (Figure 3, p=0.026). These results collectively show that environmentally relevant concentrations of PFOA had no effects on pluripotency. For the remaining experiments outlined in this paper, we utilized PFOA concentrations of 0.5, 5, and 50 μM, to model environmentally and occupationally relevant exposures.

**Figure 3:**
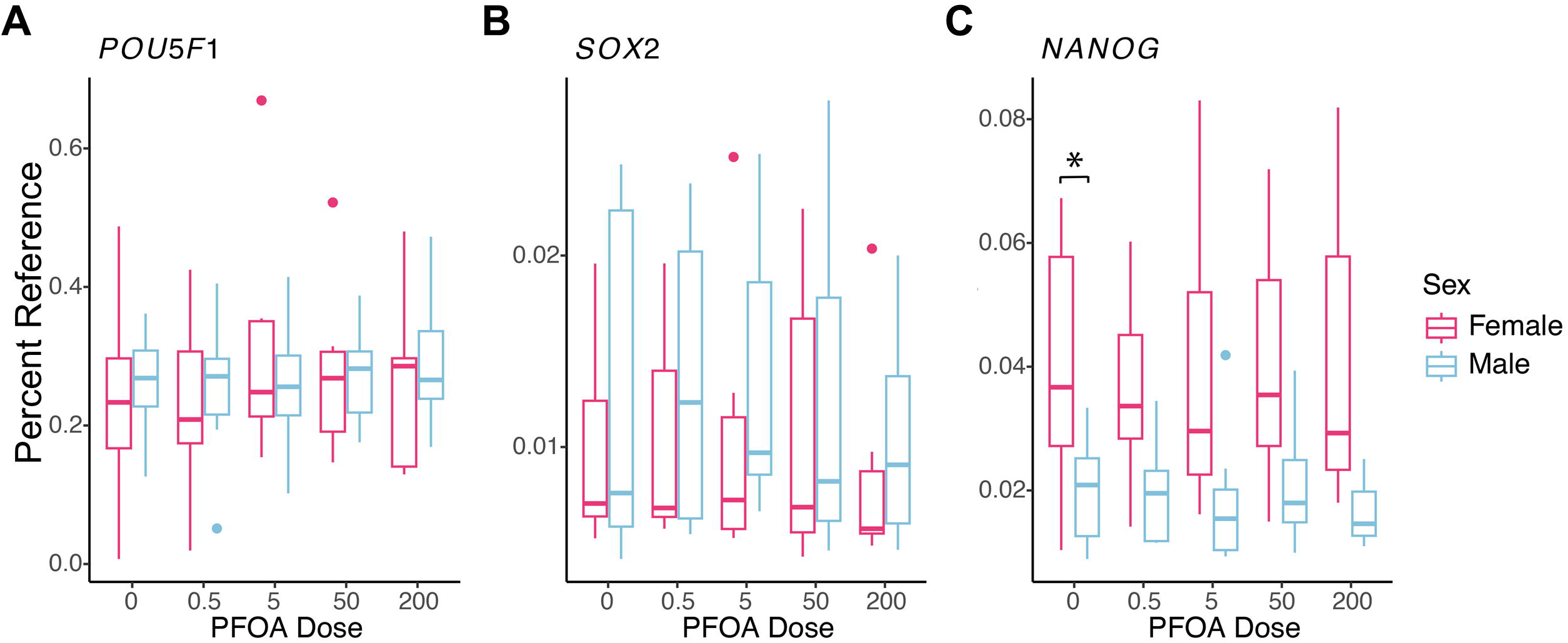
PFOA exposure and mRNA expression of pluripotency markers in hiPSCs. hiPSCs were exposed to different concentrations of PFOA for 72 hours, and pluripotency gene expression was analyzed using qRT-PCR. The relative quantification was conducted with the geometric mean of three housekeeping genes (*GAPDH*, *ATCB*, and *RPS18*) in each experiment. Data were analyzed using one-way ANOVA followed by post-hoc test. Gene expression between cell lines was analyzed at each concentration using 2-tailed, unpaired t-tests and are shown as mean ± SEM. (A) *POU5F1*, (B) *NANOG*, (C) *SOX2.* *p<0.05.

### Effects of PFOA exposure on cardiomyocyte action potential properties

We next examined whether exposure to PFOA during cardiomyocyte differentiation would impact their function. We first exposed hiPSC to 0, 0.5, 5, and 50 μM PFOA for 72 hours, followed by differentiation of the cells into ventricular cardiomyocytes (see methods) in the continued presence of PFOA. To assess whether effects of PFOA were due to developmental programming and not ongoing exposure, we stopped the exposure on day 20. Cardiomyocytes were then purified and allowed to mature on MatrixPlus plates in the absence of PFOA for a total of 10 days. The differentiation and exposure paradigm is shown in Figure 1. On day 30, we measured action potential properties in control and exposed cardiomyocytes using an optical mapping system with the voltage-sensitive probe, FluoVolt. Sample traces are shown in Figure 4A. In the female hiPSC-CM, PFOA caused a dose-dependent increase in beat frequency, which was significant at 5 and 50 μM. No significant changes in beat frequency were observed in the male hiPSC-CM (Figure 4B). Conduction velocity (Figure 4C) was unchanged in both cell lines, while action potential duration at 80 percent repolarization (APD_80_) was significantly increased with 5 μM PFOA in the male hiPSC-CM (Figure 4D); by contrast, 50 μM PFOA in female patient cells resulted in an increase in APD_80_. Lastly, we examined action potential duration triangulation (APDT), which is a measure of the interval between 30 and 90% repolarization^37^. APDT was significantly increased in female hiPSC-CM exposed to 50 μM PFOA (Figure 4E), and unchanged in the male hiPSC-CM. Thus, PFOA exposure during differentiation had significant effects on action potential properties that were dose and cell line dependent.

**Figure 4:**
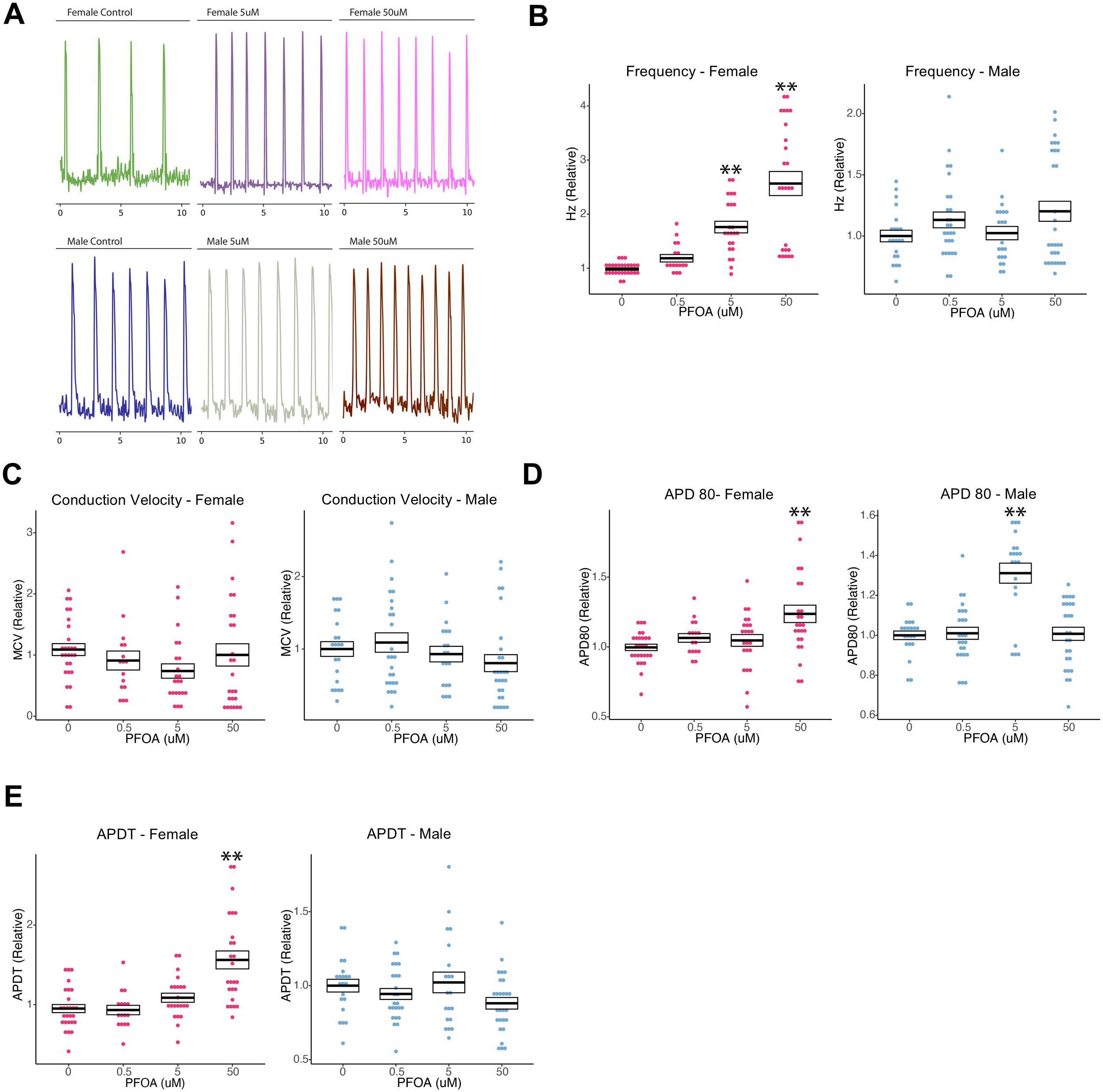
Effect of PFOA exposure during cardiac differentiation on action potential parameters of hiPSC-CMs. Each measurement was calculated relative to the mean value of the control for each experiment. Data from the female hiPSC-CM are depicted in pink, and data from the male hiPSC-CM line are shown in blue. (A) Sample 10-second time sequence plots for each condition. (B) Frequency of the action potential. (C) Conduction velocity (D) Action potential duration at 80 percent repolarization (APD 80). (E) Action potential duration triangulation (APDT). Data in panels D and E were corrected for beat frequency using Fridericia’s formula (see Methods). The data are shown as mean ± SEM. *p<0.05, **p<0.01.

### PFOA exposure and calcium dynamics

Having established that PFOA exposure impacts action potential properties, we turned our attention to calcium dynamics. Calcium plays a pivotal role in excitation-contraction coupling, the process by which cardiomyocytes couple the action potential to muscle contraction^38^. Figure 5A depicts sample optical mapping traces using the calcium-sensitive dye Calbryte 520 AM. PFOA exposure affected the frequency of calcium transients in a cell line-dependent manner. In the female hiPSC-CM we observed a dose-dependent increase in frequency, but in the male hiPSC-CM, frequency was significantly reduced with 5 μM PFOA (Figure 5B). Calcium baseline fluorescence (F0), a surrogate of diastolic calcium measurement, was significantly reduced in both cell lines by PFOA exposure. Specifically, a decrease was observed in female hiPSC-CM exposed to 5 and 50 μM, and male hiPSC-CM exposed to 5 μM PFOA (Figure 5C). Calcium transient amplitude was significantly increased with 5 μM PFOA in the female hiPSC-CM, while no significant changes were observed in the male hiPSC-CM (Figure 5D). No significant effects of PFOA exposure were observed on upstroke slope (Figure 5E) or conduction velocity (Figure 5F) in either cell line. Calcium transient duration at 80% recovery (CaTD_80_) unchanged with PFOA exposure in the female hiPSC-CM, while it was significantly increased in the male hiPSC-CM with 5 μM PFOA (Figure 5G). Calcium transient duration triangulation (CaTDT), a measure of the interval between 30 and 90% recovery, was unchanged in both cell lines (Figure 5H).

**Figure 5:**
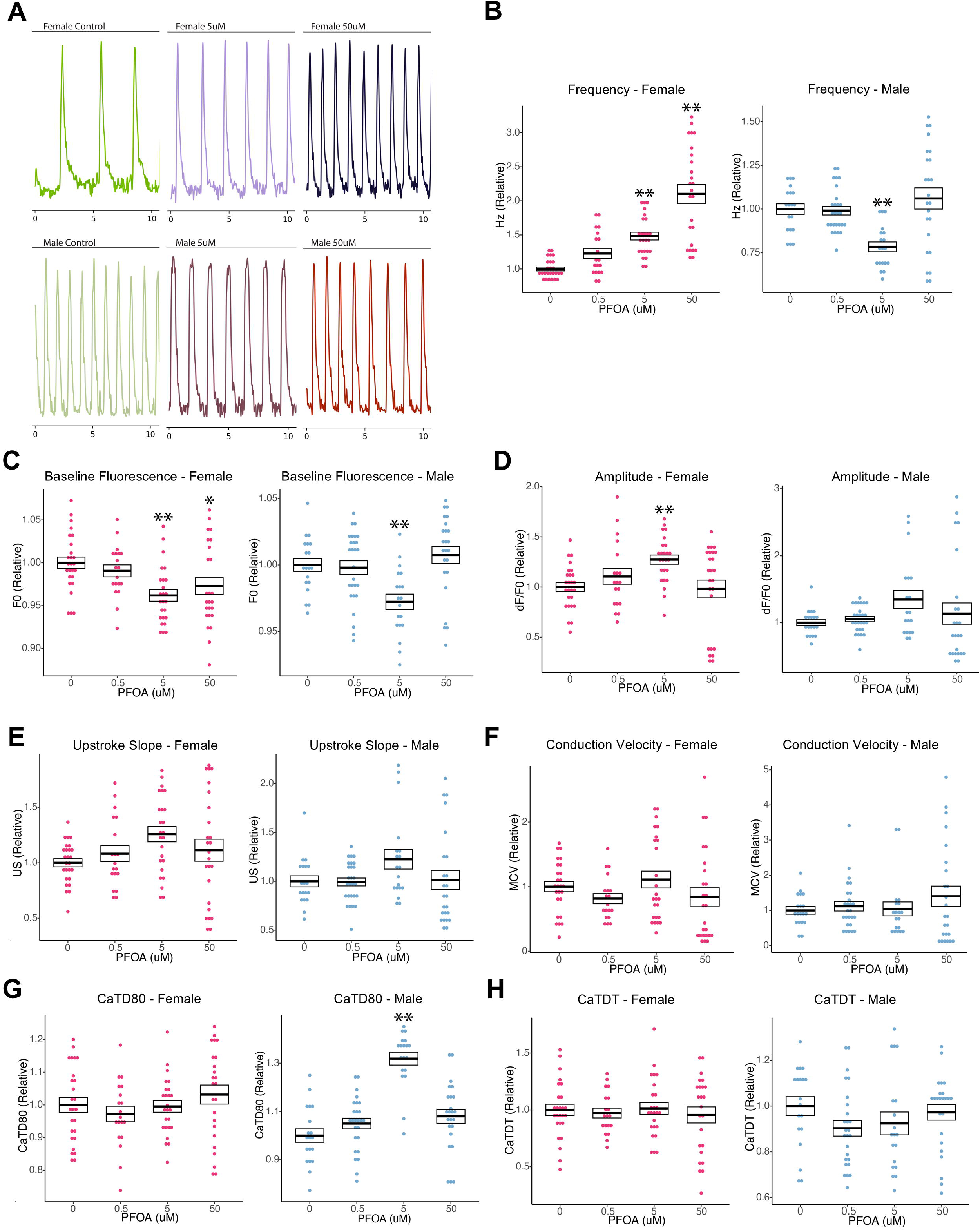
Effect of PFOA exposure during cardiac differentiation on calcium dynamics in hiPSC-CMs. Each measurement was calculated relative to the mean value of the control for each experiment. Data from the female hiPSC-CM are depicted in pink, and data from the male line are shown in blue. (A) Sample 10-second time sequence plots for each condition. (B) Frequency of the calcium transient. (C) Baseline fluorescence (F0). (D) Calcium transient amplitude. (E) Upstroke slope of calcium transient. (F) Conduction velocity. (G) Calcium transient duration at 80 percent recovery (CaTD80). (H) Calcium transient duration triangulation (CaTDT). Data in panels G and H were corrected for beat frequency using Fridericia’s formula (see Methods). The data are shown as mean ± SEM. *p<0.05, **p<0.01.

### Effects of PFOA on Calcium and Potassium Currents

We next investigated the effects of PFOA exposure on ion channel currents using whole cell voltage clamping. Specifically, we measured inward calcium channel current *I*_Ca,_ as well as the rapid delayed rectifier *I_Kr_* and inward rectifier *I_K1_* potassium channel currents. 5 μM PFOA exposure significantly reduced *I_Kr_* tail and steady-state currents in both female and male hiPSC (Figure 6A-C). Surprisingly, 50 μM PFOA did not affect *I_Kr_*compared to vehicle control, suggesting a non-monotonic dose response relationship (Figure 6A-C). PFOA minimally affected *I_K1_* (Supplemental Figure 2A-B). *I_K1_*was elevated in the male hiPSC-CM following 5 μM PFOA exposure, but only at membrane potentials hyperpolarized beyond -100 mV (Supplemental Figure 2A-B). PFOA exposure did not affect *I*_Ca,_ in male or female hiPSC-CM (Supplemental Figure 3A-B).

**Figure 6:**
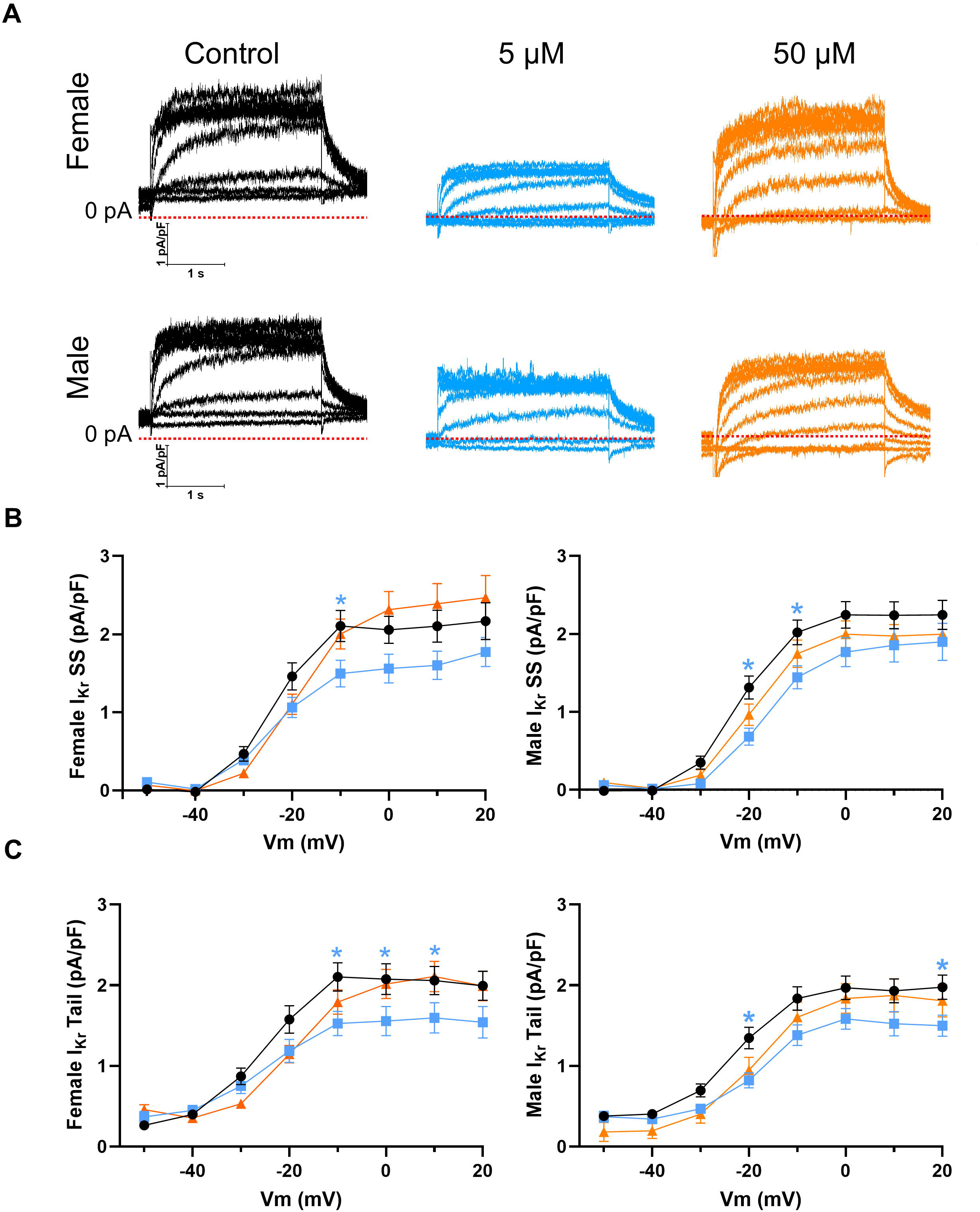
Patch clamp electrophysiology data showing effects of PFOA exposure during cardiac differentiation on *I_Kr_* currents in hiPSC-CM from the female and male donor. (A) Sample *I_Kr_* current traces for control and PFOA exposed cells. (B) Steady-state *I_kr_* measured at the end of the step pulse and (C) Tail *I_Kr_*. The data are shown as mean ± SEM. *p<0.05.

### PFOA exposure and programming of gene expression in cardiomyocytes

Having observed the effects of developmental PFOA exposure on cardiomyocyte function that differed by cell line and concentration, we sought to further identify underlying molecular targets using RNA-seq gene expression profiling. We hypothesized that we would observe changes in expression of genes related to cardiac development and muscle contraction. Principal component analysis of these data indicated that the cell line was the largest source of variation (Supplemental Figure 4). No differentially expressed genes (DEGs, genes meeting a FDR<0.05 and an absolute log2 fold change of >0.5) were identified in the female hiPSC-CM at any concentration. In the male hiPSC-CM, we identified 1950 and 22 DEGs in the 5 and 50 μM PFOA groups, respectively (Table 1 and Figure 7A-B). Among the DEGs identified in this cell line, 20 genes were differentially expressed in both exposure groups, with changes in expression occurring in the same direction (Figure 7C and Table 2). A single gene, *SPHKAP*, was up-regulated following both exposures, while all others were down-regulated. For almost all genes which changed in both exposure conditions, the change in expression was more pronounced in the 50 μM exposed cells compared to 5 μM (Table 2). The list of genes included several associated with cardiac function and development. The results of differential testing for all genes from the RNA-seq analyses are shown in the Supplemental Excel File 1.

**Figure 7:**
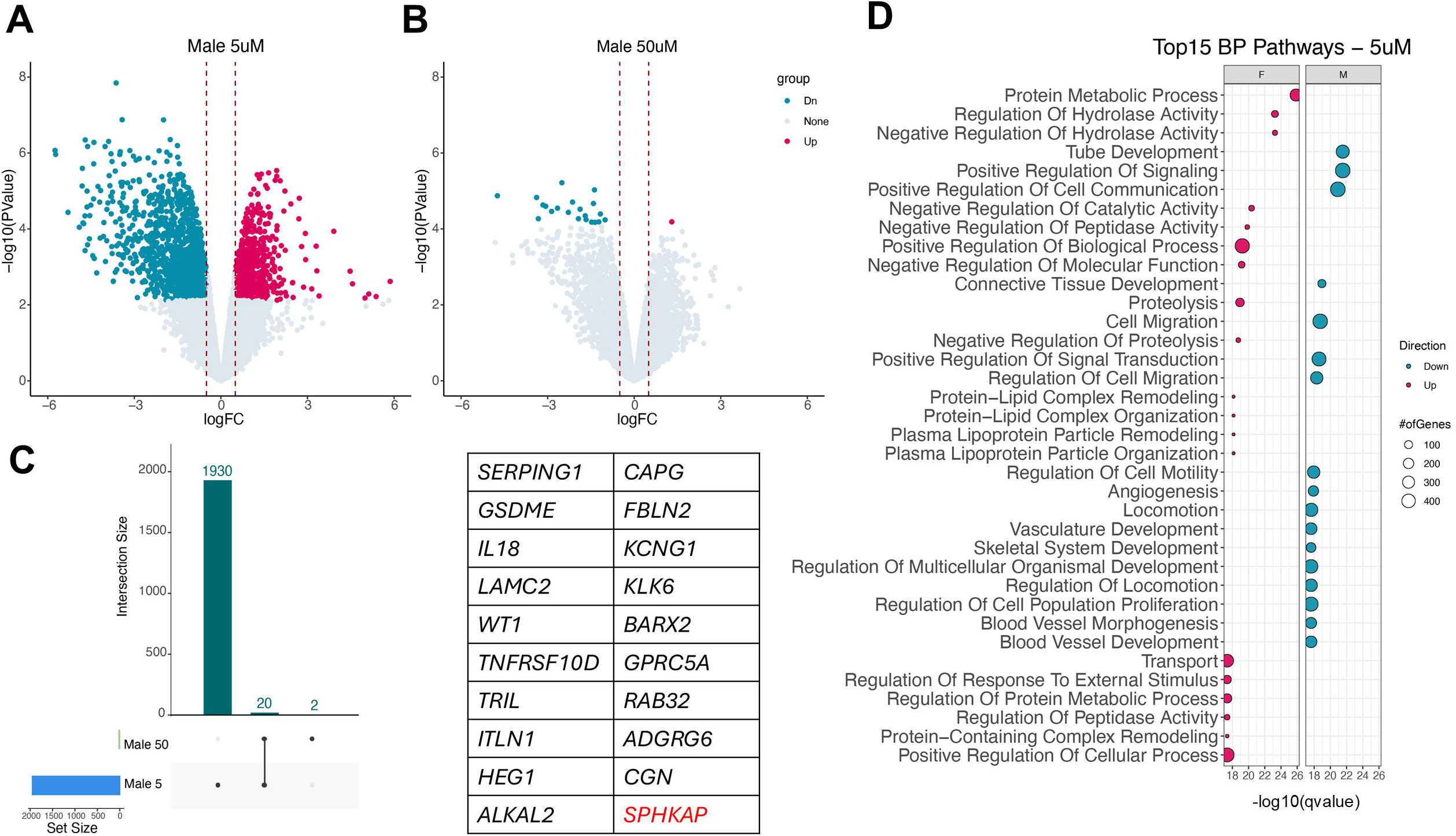
RNA-seq analysis of hiPSC-CM exposed to PFOA during differentiation. (A-B) Volcano plots showing differentially expressed genes (DEGs) for 5 and 50 μM PFOA vs. control in the male hiPSC-CM. Blue and red dots depict genes with significantly decreased and increased expression, respectively. (C) UpSet plot depicting the number of DEGs for each concentration of PFOA in the male hiPSC-CM and the degree of overlap in DEGs between concentrations. The 20 genes overlapping between 5 and 50 μM are shown on the right of the plot. *SPHKAP*, shown in red, was the only upregulated gene, while all others were down-regulated. (D) Results of the top 15 differentially expressed Gene Ontology Biological Process pathways from the female (left) and male (right) hiPSC-CM.

**Table 1:** Differentially expressed genes in ventricular hiPSC-CM exposed to PFOA during differentiation (FDR < 0.05, |LogFC| >0.5).

| | PFOA Concentration<br>(μM) | Total | Up | Down |
| --- | --- | --- | --- | --- |
| Female Patient | 0.5 | 0 | 0 | 0 |
|  | 5 | 0 | 0 | 0 |
|  | 50 | 0 | 0 | 0 |
| Male Patient | 0.5 | 0 | 0 | 0 |
|  | 5 | 1950 | 736 | 1214 |
|  | 50 | 22 | 1 | 21 |

**Table 2:** Common DEGs in male hiPSC-CM exposed to 5 and 50μM PFOA during differentiation.

| Gene | 5 μM PFOA | | 50 μM PFOA | |
| --- | --- | --- | --- | --- |
|  | Log<sub>2</sub>FC | FDR | Log<sub>2</sub>FC | FDR |
| *BARX2* | -3.38 | 0.041 | -4.62 | 0.00083 |
| *KLK6* | -3.32 | 0.042 | -4.41 | 0.0018 |
| *ITLN1* | -3.16 | 0.041 | -4.70 | 0.00083 |
| *WT1* | -3.06 | 0.042 | -3.92 | 0.0019 |
| *TRIL* | -2.86 | 0.042 | -2.98 | 0.0052 |
| *ALKAL2* | -2.65 | 0.042 | -3.80 | 0.0025 |
| *CGN* | -2.51 | 0.042 | -2.50 | 0.0019 |
| *IL18* | -2.28 | 0.042 | -3.14 | 0.0015 |
| *GPRC5A* | -2.14 | 0.042 | -3.03 | 0.013 |
| *KCNG1* | -1.93 | 0.042 | -2.87 | 0.0063 |
| *SERPING1* | -1.70 | 0.042 | -2.04 | 0.0018 |
| *GSDME* | -1.67 | 0.042 | -2.10 | 0.0021 |
| *FBLN2* | -1.48 | 0.042 | -2.29 | 0.021 |
| *CAPG* | -1.41 | 0.042 | -1.49 | 0.0040 |
| *ADGRG6* | -1.38 | 0.042 | -1.70 | 0.0042 |
| *LAMC2* | -1.36 | 0.042 | -1.18 | 0.010 |
| *RAB32* | -1.22 | 0.042 | -1.82 | 0.0042 |
| *HEG1* | -1.17 | 0.042 | -1.42 | 0.0023 |
| *TNFRSF10D* | -1.0084 | 0.042 | -1.41 | 0.0034 |
| *SPHKAP* | 1.30 | 0.042 | 1.64 | 0.0021 |

### PFOA has concentration and cell line-dependent effects on gene pathways

To gain further insight into transcriptional pathways regulated by PFOA, we performed gene set enrichment analysis. We found significantly enriched pathways in both cell lines, with differences based on PFOA concentration and cell line. Full lists of pathways are found in the Supplemental Excel File 2. Figure 7D shows the top pathways (most significant when ranked by FDR) in each cell line for 5 μM PFOA. Notably, the top pathways were distinct between cell lines. In the female hiPSC-CM, we observed upregulation of pathways related to protein metabolism, hydrolase activity, and lipoprotein remodeling. In the male hiPSC-CM, the top pathways were all downregulated and were related to development and cell migration (Figure 7D). Given that the functional effects of PFOA were highly dose-dependent, we examined the extent of overlap in regulated pathways across PFOA concentrations. In the female cell line, only a single pathway, “Regulation of System Process”, overlapped across all 3 PFOA concentrations. 57 common pathways overlapped between the 5 and 50 μM concentrations, which were primarily related to lipid metabolism (Supplemental Figure 5A). In the male donor hiPSC-CM, a total of 89 pathways were common across all 3 concentrations of PFOA (Supplemental Figure 5B). Interestingly, for the lowest PFOA concentration 0.5 μM, the pathways changed in the opposite direction compared to the 5 and 50 μM concentrations (Supplemental Figure 5B). Pathways related to muscle contraction, organ development, and calcium ion homeostasis were differentially regulated across all 3 PFOA concentrations (Supplemental Figure 5B).

### Effect of PFOA exposure on expression of calcium handling genes

Since we observed effects of PFOA on calcium dynamics (Figure 5) we next utilized the RNA-seq data to examine whether PFOA exposure impacted expression of genes involved in calcium handling. We interrogated expression of the L-type calcium channel (*CACNA1C*), the ryanodine receptor (*RYR2*), the SERCA2A calcium ATPase (*ATP2A2*), the sodium-calcium exchanger (*SLC8A1*), and phospholamban (*PLN*). None of these genes met the FDR criteria for differential expression, so we examined the RPKMs for each gene individually. Notably, we found lower baseline (control groups) expression of all 5 genes in the control male hiPSC-CM compared to the female cells, which was statistically significant for *ATP2A2* and *SLC8A1* (p<0.05, Supplemental Figure 6). PFOA exposure had no significant effects on gene expression in the female hiPSC-CM (Figure 8A-E). In the male hiPSC-CM, however, we found that 5 and 50 μM PFOA resulted in increased expression of all 5 genes, with changes that were statistically significant for *CACNA1C, RYR2,* and *SLC8A1.* In contrast, although not statistically significant, 0.5 μM PFOA exposure resulted in lower expression of these genes in both cell lines (Figure 8A-E).

**Figure 8.**
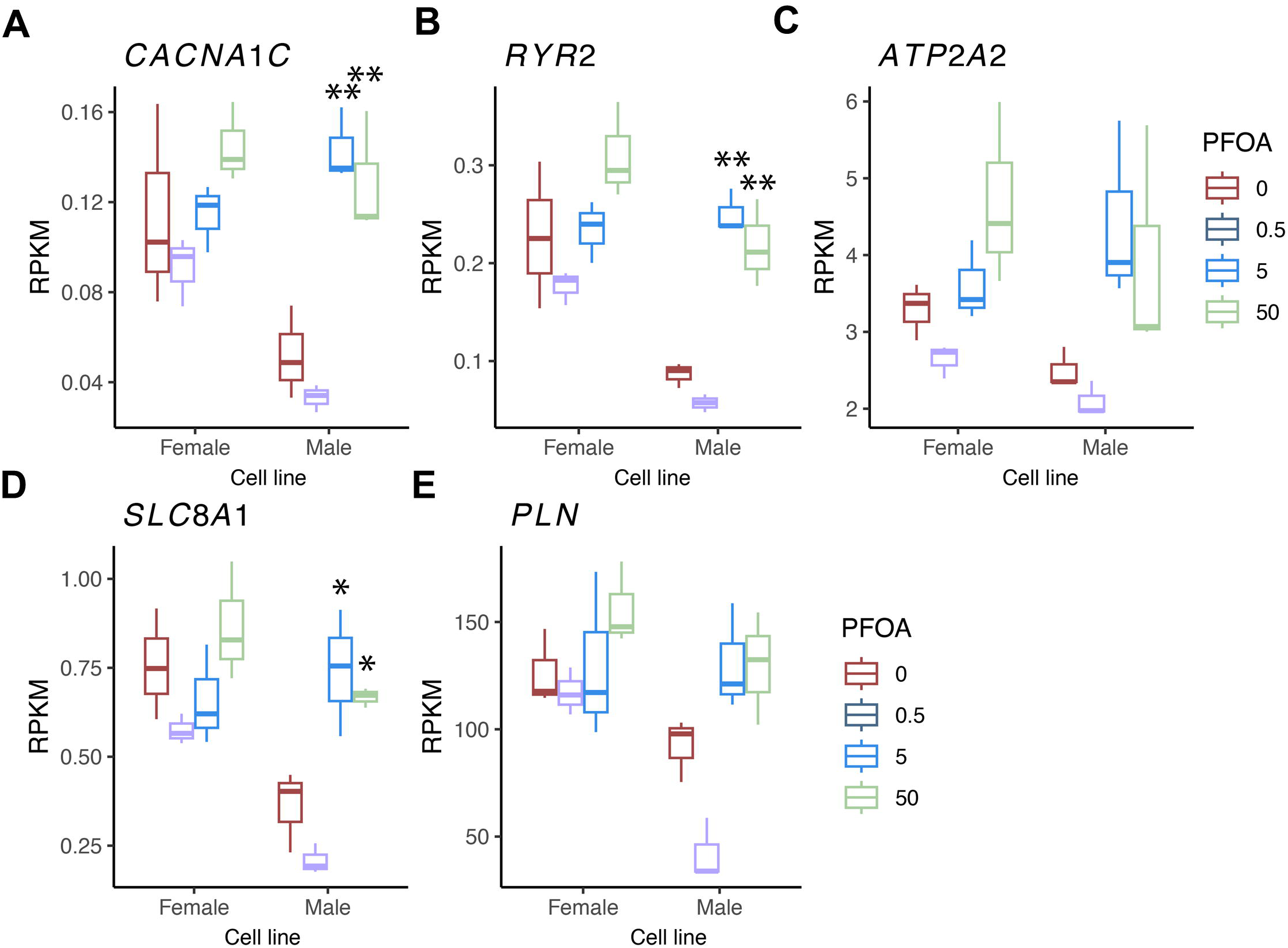

## Discussion

Exposure to per- and polyfluoroalkyl substances is associated with various cardiovascular diseases, but effects of these chemicals on the developing human heart are poorly understood. Using an *in vitro* model of cardiac differentiation, in this study we examined the functional and transcriptomic effects of one of the most common PFAS, perfluorooctanoic acid, on ventricular cardiomyocytes. To our knowledge, this is the first study in differentiating human stem cells which demonstrates that PFOA exposure has impacts on cardiomyocytes that persist long after cessation of exposure.

### Pluripotency and Proliferation

Our data show that PFOA exposure had no significant effects on expression of pluripotency markers, even at the highest concentrations. Two studies in human pluripotent stem cells revealed similar findings. Recent work in human embryonic stem cells (hESCs) using perfluorooctane sulfonic acid (PFOS), another C8 PFAS, showed reduced expression of pluripotency markers only at the highest concentration, 50 μM^39^. A second study in hiPSC embryoid body cultures showed no significant effects on expression of pluripotency markers *OCT4* or *NANOG* with PFOA concentrations of up to 100 μM^40^. With regard to proliferation, we observed a significant reduction only at the highest concentration (200 μM) of PFOA tested, in line with inhibitory effects of PFOS at similarly high concentrations^39^. Notably, lower concentrations of PFOA led to modest *increases* in proliferation, suggesting a potential non-monotonic response. Such responses are increasingly recognized in PFAS toxicology and may reflect adaptive or compensatory cellular processes at lower exposure levels^41^. Importantly, these findings may indicate that the functional and transcriptional changes observed in differentiated cardiomyocytes are unlikely to be secondary to overt toxicity but rather reflect more subtle alterations in developmental programming. These data collectively suggest that human-relevant concentrations of these long chain PFAS are not acutely cytotoxic to stem cells *in vitro*.

### Action potential and calcium dynamics

Our work demonstrated that PFOA exposure during differentiation impacted action potential and calcium dynamics in the cardiomyocytes, with effects that persisted for several days after cessation of exposure. Whether these effects diminish beyond the time frame of this study is an important area for further investigation. Reductions in beat rate and pulsation amplitude have been reported in H9 hESC-cardiomyocytes exposed to PFOS during differentiation^39^, but persistent effects after cessation of PFOA exposure were not examined. Likewise, acute effects of 56 different PFAS, including PFOA, on hiPSC-CM from 16 different donors were examined, and showed effects on calcium dynamics and cytotoxicity that varied markedly between individual PFAS as well as patient donors^42^. Our findings thus add to a limited but growing body of evidence that PFAS have persistent impacts on cardiomyocyte function, and are in keeping with human epidemiologic^14–16,43^ and animal ^20,21^ data showing adverse cardiovascular effects of these chemicals.

Critically, the persistence of these functional alterations after removal of exposure suggests that PFOA may induce developmental reprogramming of cardiomyocyte physiology, rather than transient functional disruption. This distinction is central to understanding PFAS toxicity, as it implies that brief exposures during critical windows of differentiation may have long-lasting consequences for cardiac function^44^. Such persistence is consistent with the Developmental Origins of Health and Disease (DOHaD framework and supports the hypothesis that environmental exposures during early development can shape long-term cardiovascular risk^45^.

To better understand potential mechanisms, we investigated whether the observed changes in action potential and calcium properties were due to alterations in ion channel function, including *I_Ca_*, predominantly conducted by Ca_V_1.2^46^ as well as *I_K1_* and *I_Kr_*, conducted by Kir2.1/2.3 and hERG1 potassium channels, respectively^47–49^. PFOA in the extracellular bath of hiPSC-CM was recently reported to rapidly inhibit *I_Ca_* and *I_Kr_* by disrupting channel transcript levels (< 30 mins)^50^. Our recordings demonstrate that the effects on *I_Kr_* and *I_K1_*magnitude persist more than 10 days after PFOA removal. The selective increase in *I_K1_* in male cells by PFOA may explain the absence of an effect of PFOA on male cardiomyocyte firing frequency.

Beyond confirming prior acute findings, these results indicate that PFOA exposure during differentiation leads to stable remodeling in ion channel function, particularly affecting repolarization currents^50,51^. Given the central role of *I*_Kr_ in controlling APD, these changes may contribute to increased susceptibility to arrhythmogenic events, especially under stress conditions not examined in this study.

Our data also suggest that PFOA impacts the expression and function of calcium handling machinery. In response to an action potential, L-type calcium channels (*Ca_V_1.2*/*CACNA1C*) open and allow calcium to enter the cardiomyocyte. This influx of calcium activates ryanodine receptors (*RYR2*) to release calcium from the sarcoplasmic reticulum, in a process known as calcium-induced calcium release^38^. This increased cytoplasmic calcium binds to myofilaments to initiate muscle contraction^38^. At the end of contraction, calcium is re-sequestered in the SR via the SR ATP-dependent calcium pump (SERCA2A/*ATP2A2*)^38^. The sodium-calcium exchanger (*SLC8A1*) further regulates cytoplasmic calcium. Notably, we found that PFOA exposure resulted in a significant increase in expression of several genes in this pathway, including *CACNA1C, RYR2,* and *SLC8A1*, with a trend toward increased expression of *ATP2A2* in the male hiPSC-CM, and a trend toward increased expression of all 4 genes (*CACNA1C, RYR2, ATP2A2,* and *SLC8A1*) in the female hiPSC-CM. Each of these genes is developmentally regulated^52–55^, suggesting that PFOA may impact their transcriptional regulation through effects on the epigenome and/or transcriptional machinery. Likewise, calcium handling and excitation-contraction coupling are strongly regulated by posttranslational mechanisms^38^; thus future studies examining these mechanisms are warranted. Taken together, these findings suggest that PFOA exposure disrupts multiple interconnected components of excitation-contraction coupling, resulting in coordinated changes in both electrical activity and calcium signaling. This integrated disruption may represent a key mechanistic pathway linking PFAS exposure to impaired cardiac function.

### Gene expression changes following PFOA exposure

Gene expression profiling of cells exposed to PFOA revealed effects on genes and pathways that were distinct between cell lines. Many of the observed changes in gene expression were consistent with established mechanisms of PFAS action, as well as with the functional changes we observed. We found many DEGs in the male hiPSC-CM, with several genes that were common across the two higher PFOA concentrations. Importantly, the presence of a large number of DEGs at 5 μM but substantially fewer at 50 μM suggests a non-monotonic transcriptional response, reinforcing the idea that intermediate, environmentally relevant concentrations may exert the more pronounced biological effects. This observation challenges traditional dose-response assumptions and highlights the need for careful consideration of exposure levels in toxicological studies.

For example, in male hiPSC-CM we observed up-regulation of *SPHKAP* at both 5 and 50 μM PFOA. *SPHKAP* encodes an A-kinase anchoring protein which has been shown to interact with protein kinase A (PKA) and influence adrenergic signaling and calcium dynamics in the heart and is dysregulated in heart failure^56^. As our studies were conducted in cardiomyocytes without stress, investigating how exposure to PFOA and other PFAS impacts adrenergic signaling in cardiomyocytes is an important future direction. Pathway analysis also demonstrated distinct effects between the 2 cell lines. In the female hiPSC-CM, it is notable that we observed enrichment for pathways related to lipids and triglyceride metabolism (Figure 7D and Figure S5), given the well-established role for PFAS in dysregulation of cholesterol homeostasis^17,57^. In the male hiPSC-CM, developmental pathways and those related to heart contraction were significantly altered, consistent with the observed changes in voltage and calcium dynamics. These findings highlight that PFOA exposure leads to divergent biological responses depending on the cellular context, with metabolic pathways predominating in female hiPSC-CMs and developmental/electrophysiological pathways in male hiPSC-CMs. Such divergence suggests that baseline cellular state and intrinsic gene expression differences may shape susceptibility to PFAS exposure^42,58–60^.

Likewise, we found significant changes in pathways related to immune system function and cytokine production. PFAS effects on immune cells have been widely reported^9^, but less is known about their effects on cytokine signaling in the cardiovascular system. However, recent work suggests that PFAS may also exert toxic effects on the heart through inflammatory mechanisms^61^. Although the gene expression data do not provide full mechanistic insight into the effects of PFOA, they nevertheless demonstrate significant dysregulation of transcriptional programs, particularly in the male hiPSC-CM, even days after cessation of exposure. Importantly, the presence of both shared and dose-specific pathways suggests that PFOA may engage multiple overlapping and dose-dependent molecular mechanisms, rather than a single linear pathway^40,62,63^.

### Summary, Limitations, and Public Health Implications

Exposure to environmental contaminants may have adverse cardiovascular effects that persist across the life course^64^. Using a New Approach Methodology focused on cardiovascular development, this study addresses several important knowledge gaps related to the effects of PFAS on cardiovascular health, demonstrating that PFOA exposure during cardiomyocyte differentiation has effects on transcription and function which persist after cessation of exposure and are cell line dependent. The observed effects of PFOA on voltage and calcium dynamics are notable, given that both prolongation and shortening of action potential duration can precipitate cardiac arrhythmias^65^, and several cardiovascular diseases, including heart failure and cardiomyopathies^66^, are characterized by dysregulation of calcium handling. Notably, these studies were conducted at baseline, without additional stressors and it will be important to investigate whether PFOA and other PFAS impact the ability of cardiomyocytes to respond to physiologic and pathophysiologic stress. Our data demonstrating marked inter-individual variability in the effects of PFOA are consistent with recent findings^42^. An important limitation of this study is the use of 2 cell lines, from a single male and female donor. It is imperative that we better understand the sex-specific effects of PFAS on the heart, but to fully understand whether the differences observed between cell lines are due to sex or some other variable, additional cell lines and/or isogenic cells^67^ are necessary. Recent “village in a dish” approaches which pool several hiPSC lines show promise for understanding effects of exposures on a larger population of patient-derived cells^68^. The mechanism by which effects of PFOA exposure persist beyond the exposure period is currently unclear. PFOS (another C8 PFAS) exposure was shown to alter DNA methylation in differentiating cardiomyocytes^39^, suggesting that some PFAS may have persistent functional effects through programming of the epigenome. A second putative mechanism involves incorporation of PFOA into the lipid bilayer of the cell membrane, altering membrane fluidity and ion channel function. This was recently reported to occur in human platelets^69^ as well as rat hearts^70^. Together, these findings indicate a model in which developmental exposure to PFOA induces persistent reprogramming of cardiomyocyte function through combined effects on transcriptional regulation, ion channel activity, and calcium handling. PFOA is one of a class of chemicals comprised of thousands to millions of structures, and the health effects of most PFAS are poorly defined^71^. Our findings highlight the urgent need for research into effects of other, structurally diverse PFAS on stem cell and cardiomyocyte biology. Lastly, this *in vitro* model cannot capture the effects of PFAS on cardiac structure, nor can it shed light on cardiovascular effects that are secondary to neurohumoral, inflammatory, and other mechanisms. Additional studies in more complex models such as human heart organoids^72^ and microphysiological systems are an important future direction.

## Acknowledgements

Funding for this work was provided by a NIEHS Transition to Independent Environmental Health Researcher (TIEHR) K01 (ES032048), a NIEHS Revolutionizing Innovative, Visionary Environmental Health Research (RIVER) R35 (ES031686), the Michigan Center on Lifestage Environmental Exposures and Disease P30 Center (M-LEEaD), NIEHS T32 (ES007062), NIEHS R01 (ES028802), NHLBI R01 (HL171039), and the Michigan Biological Research Initiative on Sex Differences in Cardiovascular Disease (M-BRISC).

Data Sharing: All sequencing data will be uploaded to GEO upon acceptance of this manuscript.

## Supplemental Figure Legends

**Figure S1:** QIAxcel PCR products showing expected band sizes of 254 bp for SRY (found on the Y chromosome) and 300 bp for ATL1 (found on the X chromosome).

**Figure S2:** Patch clamp electrophysiology data showing effects of PFOA exposure during cardiac differentiation on *I_K1_* currents in hiPSC-CM from the female and male donor. (A) Sample *I_K1_* current traces for control and PFOA exposed cells. (B) Current-voltage relationship for *I_k1_*. Insets show the expanded outward component of *I*_K1_. The data are shown as mean ± SEM. *p<0.05.

**Figure S3:** Patch clamp electrophysiology data showing effects of PFOA exposure during cardiac differentiation on *I_Ca_* currents in hiPSC-CM from the female and male donor. (A) Sample *I_Ca_* current traces for control and PFOA exposed cells. (B) Current-voltage relationship for *I_Ca_*. The data are shown as mean ± SEM. *p<0.05.

**Figure S4:** PCA plots of RNA-seq data showing the clustering of each sample on the basis of cell line (A,C) and concentration (B,D).

**Figure S5:** Pathway analysis results in the female- and male hiPSC-CM (A and B, respectively), showing overlap in enriched pathways between PFOA concentrations. In the female hiPSC-CM, only a single pathway overlapped between all 3 concentrations, so only pathways for 5 and 50 μM PFOA are shown. The size of each circle reflects the number of genes in each pathway, and the color indicates direction (red and blue for up and down-regulated, respectively).

**Figure S6**: Gene expression data for calcium handling genes in control hiPSC-CM from each donor (female in red, and male in blue). Data reflect normalized RNA-seq read counts, and were analyzed by unpaired t-test. * Denotes p<0.05.

