## Supplemental Figures for "Acute Exposure to Perfluorooctanoic Acid (PFOA) During Cardiomyogenesis disrupts Transcriptional and Electrophysiological Profiles in Differentiated Myocytes"

### Slide 1
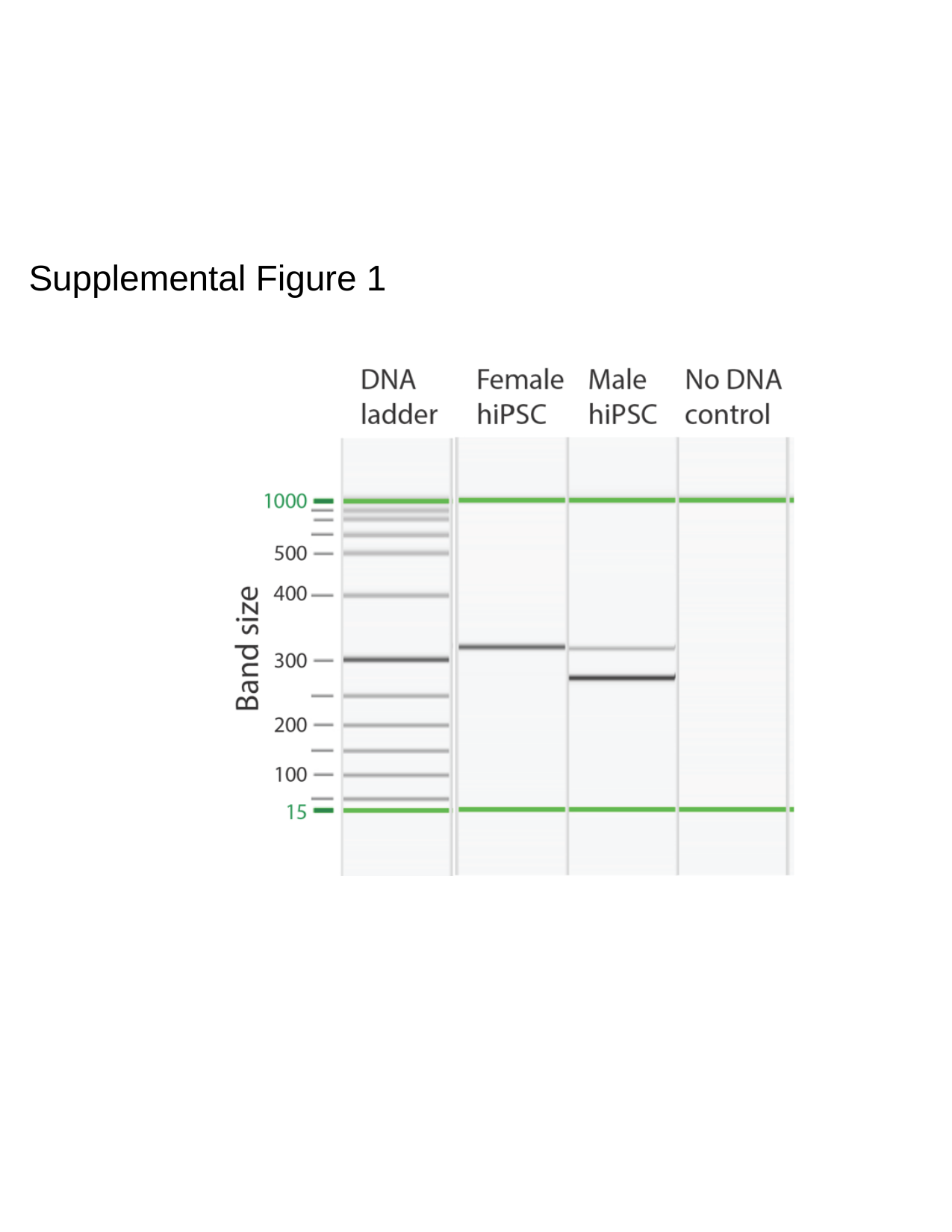

Supplemental Figure 1

### Slide 2
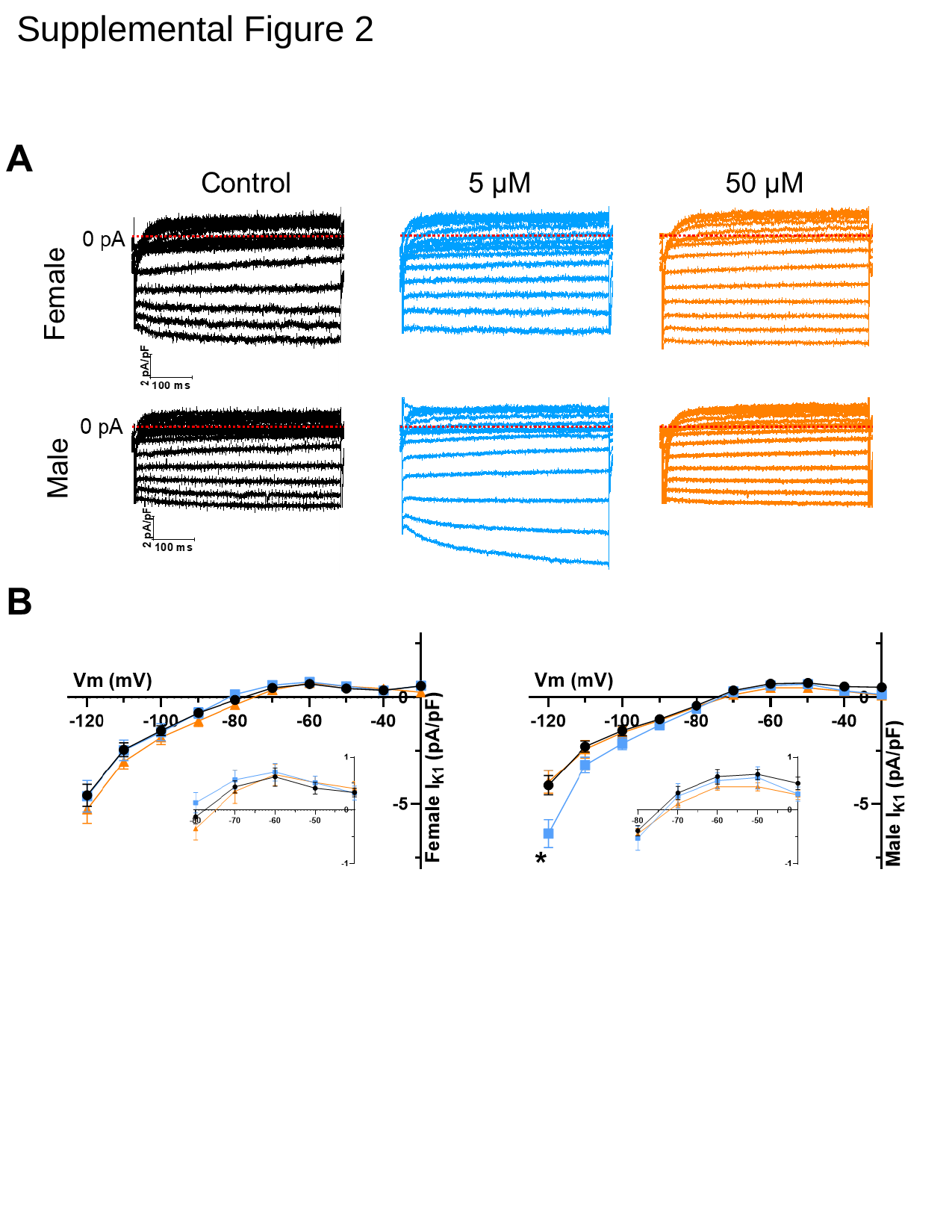

Supplemental Figure 2

### Slide 3
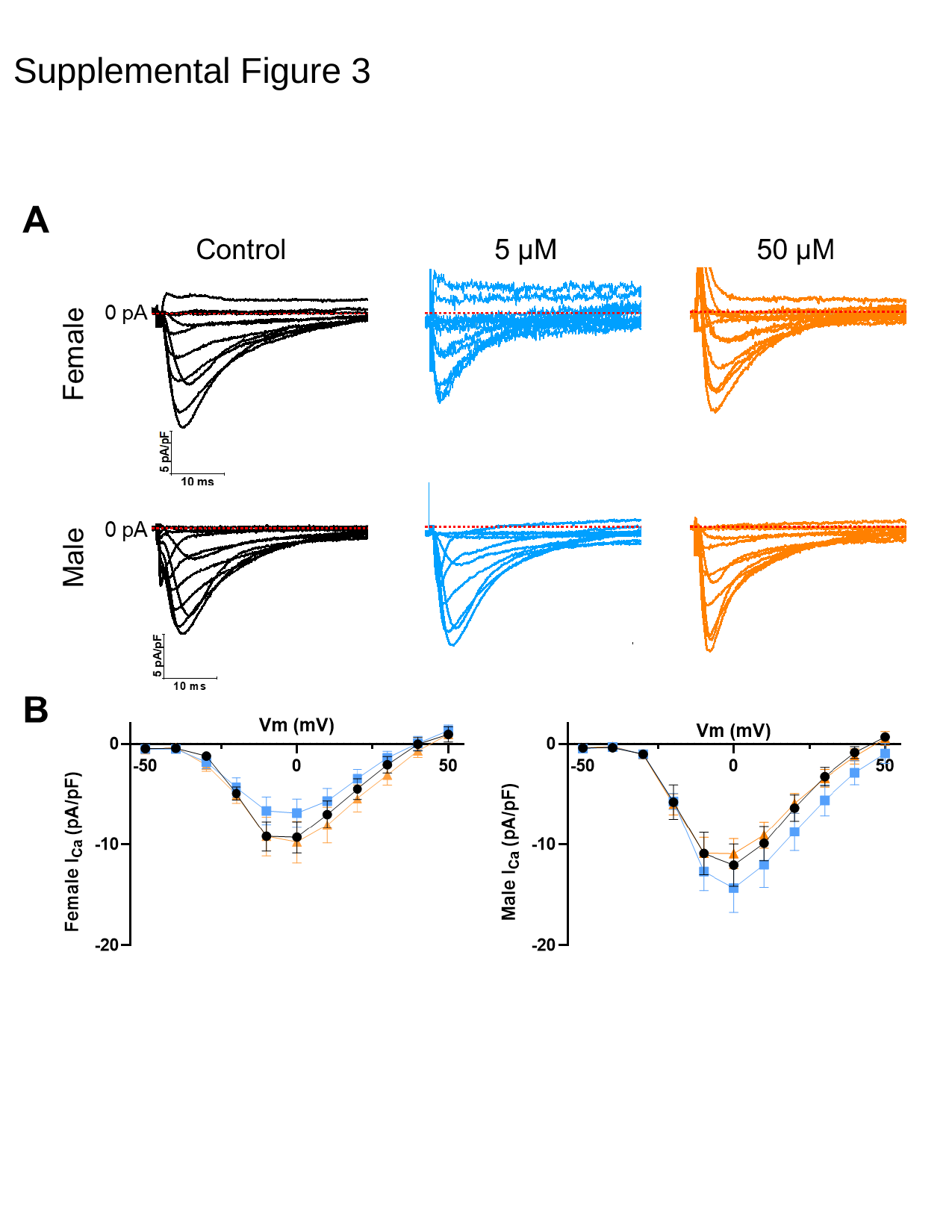

Supplemental Figure 3

### Slide 4
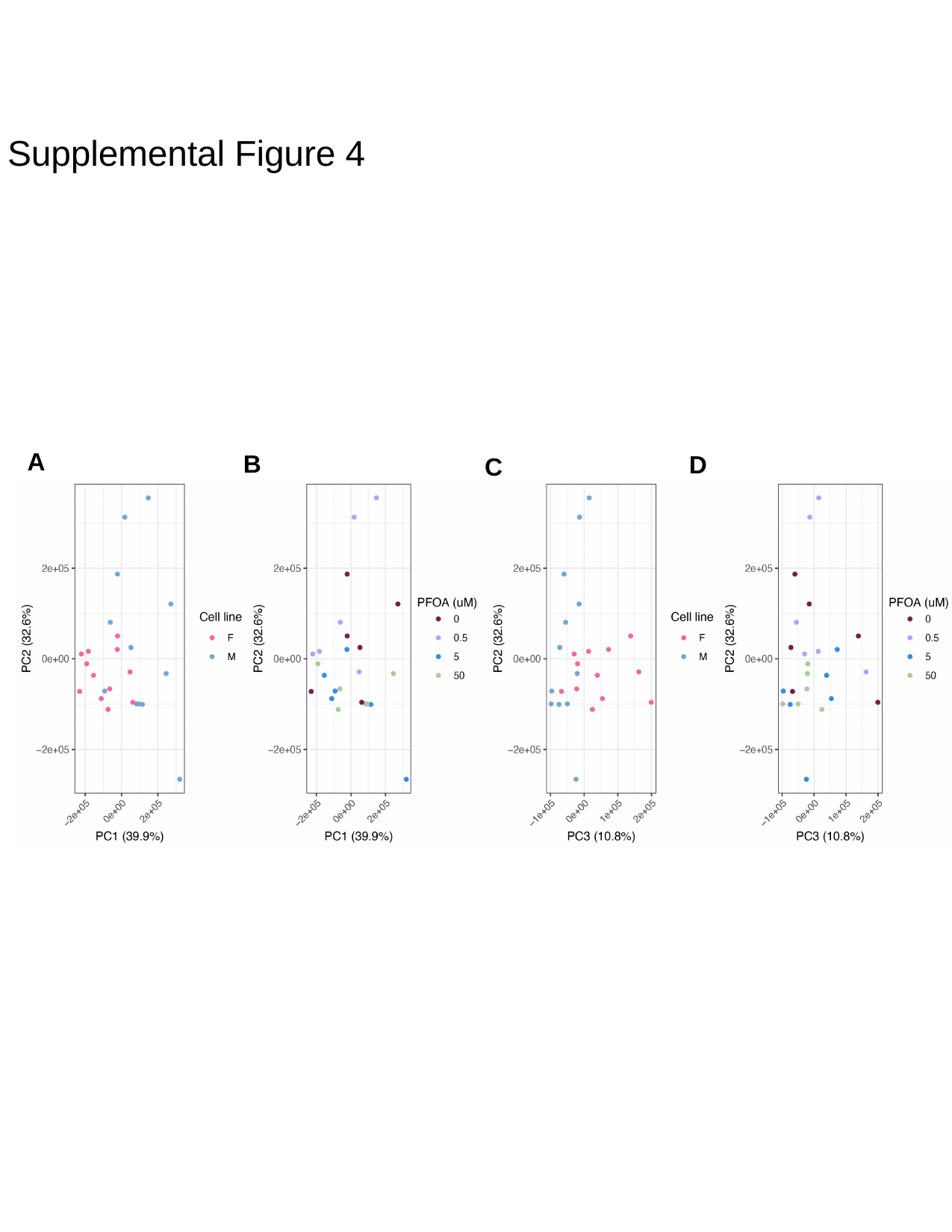

Supplemental Figure 4
A
B
D
C

### Slide 5
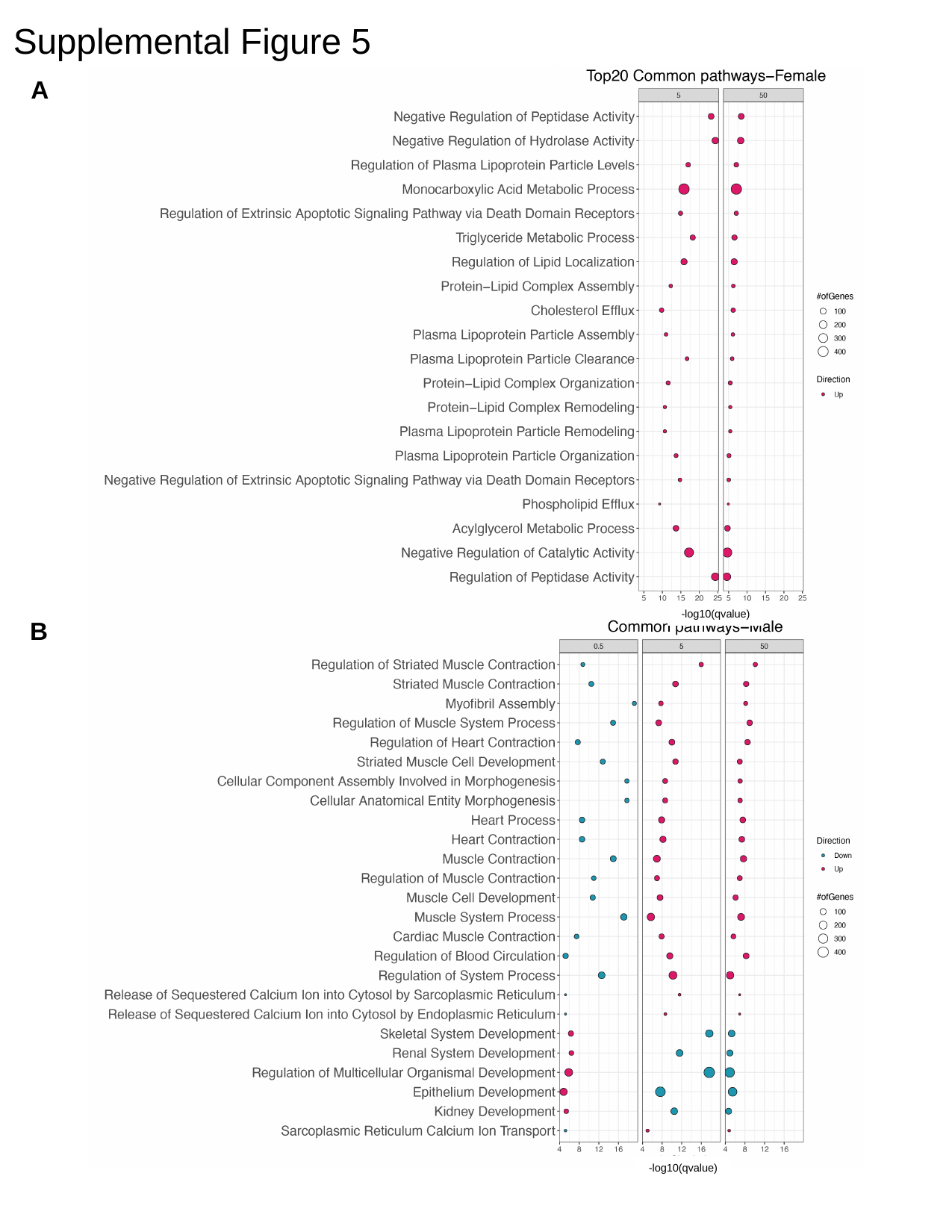

Supplemental Figure 5
A
-log10(qvalue)
B
-log10(qvalue)

### Slide 6
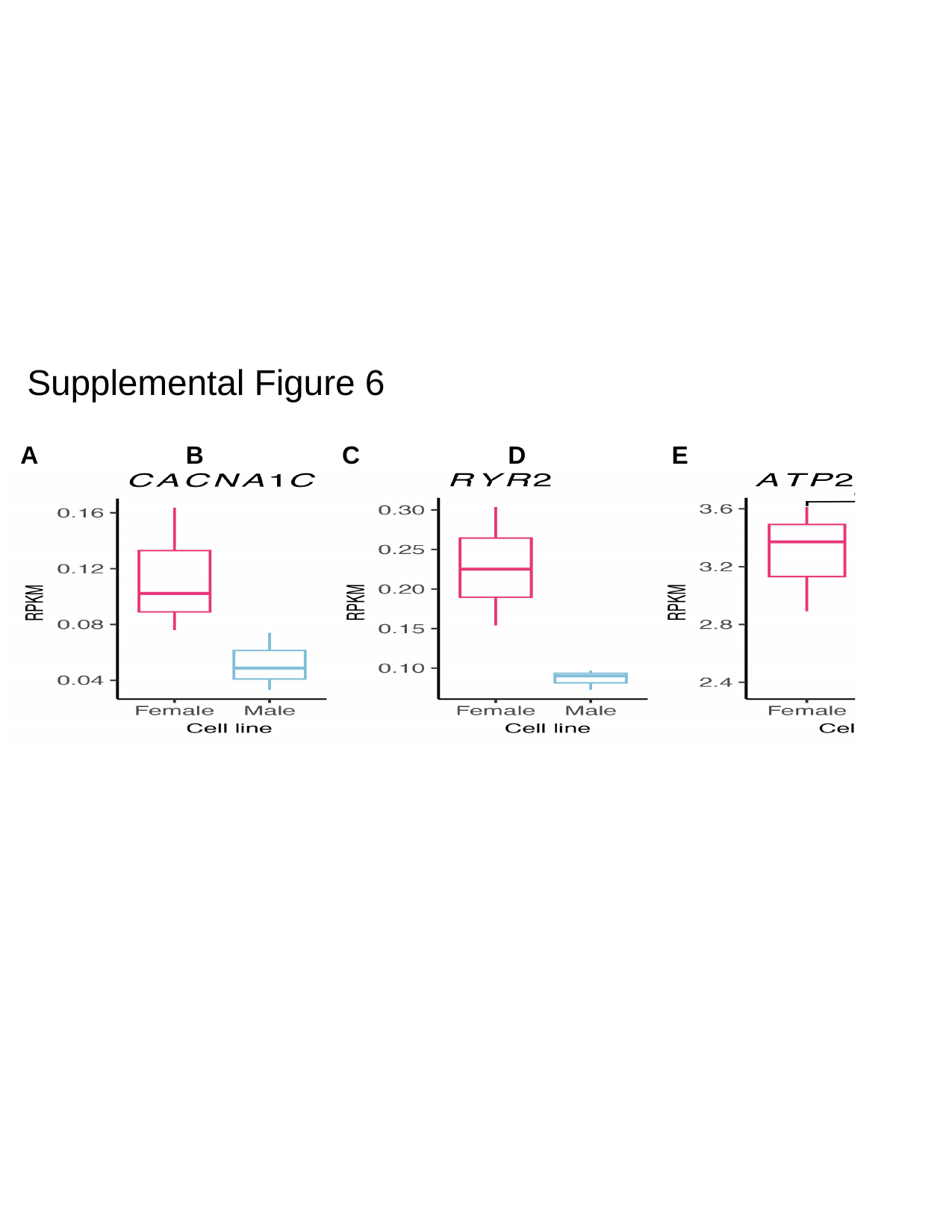

Supplemental Figure 6
A
C
D
E
B
